# Mitochondrial morphodynamics are modulated by physiological range of temperature and influence host cell outcomes during influenza infection

**DOI:** 10.1101/2024.09.06.611682

**Authors:** Grant J. McLoughlin, Andrew Pekosz

## Abstract

Influenza viruses replicate in both the cooler, upper portions of the airway and the warmer, lower portions of the respiratory tract. This study investigates how physiological ranges of temperature, specifically 33°C and 37°C, impact host cell biology and how temperature-dependent differences in host cells influence outcomes during influenza A virus infection. This study prioritizes describing mitochondrial networks due to their importance in maintaining cellular homeostasis and mediating immune responses to viral infection. The temperature at which cells are incubated significantly influences mitochondrial network morphology and mitochondrial function. Moreover, temperature-dependent changes to mitochondrial networks prior to infection result in temperature-specific changes to host cell outcomes during infection. These findings indicate that mitochondrial structure alone can modulate host cell outcomes during viral infection and that both the form and function of mitochondria directly impact influenza A virus production. While not all mitochondrial processes were shown to be affected by temperature or infection, these results highlight the importance of using physiologically relevant temperatures in respiratory pathogen research and elucidate how mitochondrial dynamics contribute to host cell outcomes during influenza A virus infection.

**Importance:** Respiratory viruses infect the upper and lower respiratory tract but rarely is the impact of physiological ranges of temperature (33°C to 37°C) considered. Mitochondria are central mediators of numerous physiological pathways, and their functions are often modified by virus infection. Physiological ranges of temperature can alter mitochondrial form and function, which is further impacted by virus infection. The study sheds light on how temperature can impact mitochondrial form and function in concert with virus infection.

## Introduction

The human respiratory tract is a physiological gradient of epithelial cell types, humidity, pressure, and temperature. During respiration, air inhaled from the ambient environment is gradually warmed as it travels from the nasal cavity to the alveoli (1). This creates distinct temperature microenvironments in the respiratory tract with the upper respiratory tract maintaining an average temperature of 33°C and the lower respiratory tract maintaining an average temperature of 37°C (2, 3). Influenza A virus (IAV) affects the upper and lower respiratory tract (4), however, most in vitro studies are performed at 37°C. While early studies indicate that physiological temperature impacts respiratory virus replication, pathogenicity, and host immune responses (5–7), but the underlying temperature-induced changes to host cell biology driving differences in viral and host cell outcomes remain unclear.

Host cells detect IAV via toll-like receptors 3 and 7 in endosomes during viral entry, and through RIG-I as the virus replicates (8). Activation of these pathogen recognition receptors induces a drastic shift in host cell transcriptional profiles to generate early type I interferon responses which help control virus replication. Additionally, IAV infection can directly cause significant alterations in host cells (9) via the viral hijacking of host cell machinery which leads to changes in cellular respiration, mitochondrial function, and the eventual induction of cell death pathways including apoptosis (10). These changes can result in cellular stress and damage, contributing to the overall pathology of the infection. Furthermore, physiological ranges of temperature can have drastic effects on the baseline transcriptional pattern of human nasal epithelial cell (hNEC) cultures and can lead to temperature dependent induction of gene expression after infection with IAV or SARS-CoV-2 (6). Antiviral related genes are activated at both temperatures, but certain antiviral responses are stronger at 37°C compared to 33°C (6).

In addition to their essential role in energy production, mitochondria are central mediators of many cellular processes including antiviral innate immune responses, programmed cell death, and ion homeostasis (11). The structure and function of mitochondrial networks, including mitochondrial fission and fusion, are particularly important for maintaining cellular health as changes in these processes can influence mitochondrial quality control, apoptosis, and metabolic adaptation during stress responses (12). Focusing on mitochondria as key mediators of host-pathogen interactions is an effective strategy to better understand how cells respond to viral infections and how virus infection impacts host cell dynamics.

In this study, changes in the structure and function of mitochondrial networks in kidney epithelial cells incubated at different physiological temperatures were analyzed to determine the effect of temperature on mitochondria, a key signaling organelle during IAV infection. Changes to mitochondrial networks and host cell outcomes during IAV infection were then analyzed at different physiological temperatures. The data provide the first known report of temperature-induced changes to mitochondrial dynamics in the context of IAV infection and describe temperature specific host cell outcomes of infection. Mitochondrial structure, but not mitochondrial function, is important for mediating temperature-specific impacts of infection. Additionally, some mitochondrial-related host cell outcomes can be altered by changing mitochondrial morphodynamics while others occur regardless of network properties.

## Materials and Methods

### Cell Culture

Madin-Darby canine kidney cells (provided by Dr. Scott Hensley, University of Pennsylvania) were maintained in complete media (DMEM supplemented with 10% fetal bovine serum, 100 units/ml penicillin/streptomycin (Life Technologies) and 2 mM Glutamax (Gibco)) at 5% CO2. Prior to all experiments, cells cultured at 37°C were split into independent, but paired flasks and incubated at 37°C and 33°C for 96 hours.

### Viruses and Virus Infections

Viruses were isolated from influenza patients in Baltimore, Maryland during the 2017-2018 and 2018–2019 influenza seasons. A/Baltimore/R0227/2017 (H3N2) and A/Baltimore/R0675/2019 (H1N1) were used for this study. Prior to infection cells were washed twice with PBS plus 100 mg/L each of anhydrous calcium chloride and magnesium chloride hexahydrate (PBS +/+). All infections used a multiplicity of infection (MOI) of 5 in infection media (DMEM supplemented with 0.3% BSA (Sigma), 100 units/ml pen/strep (Life Technologies), 2 mM Glutamax (Gibco), and 5 µg/ml N-acetyl trypsin (Sigma-Aldrich) for one hour in the minimal volume acceptable for coverage of the well or chamber. After one hour the cells were washed twice with complete media and incubated in complete media for the remainder of the experiment.

### Mdivi-1 and CoQ10 Treatments

Cells were incubated at 33°C or 37°C in complete media containing 25 µM Mdivi-1 (Tocris) in DMSO or 500 µM Conezyme Q10 (Sigma) in Pluronic F-127 (ThermoFisher) for 12 hours prior to infection. Mdivi-1 and CoQ10 treatments were removed for the infection and resumed after infection.

### TCID50 Assay

IAV infectious titers were determined by TCID_50_. MDCK-SIAT cells were seeded in 96-well plates and grown to confluence. Prior to infection cells were washed twice with 100 µl per well of PBS +/+ and once with 180 µl of infection media per well. One-hundred-fold serial dilutions were performed on viruses and 5 µl of the dilution was added to the cell plate. Each virus dilution was added to cell plates in sextuplicate, plates were incubated for six days at 33°C before being fixed with 4% paraformaldehyde for > 3h and stained overnight with naphthol blue-black. 50% tissue culture infectious dose was calculated as previously described (13, 14).

### Immunofluorescence Microscopy

For immunofluorescence microscopy, cells were plated on 18-mm glass coverslips (Fisher) or µ-Slide 8 well imaging chambers (Ibidi) and fixed with 4% paraformaldehyde (Electron Microscopy Sciences) in PBS for 15 min at room temperature. Cells were then washed once with PBS, permeabilized with PBS/0.3% Triton X-100 for 10 min at room temperature and then blocked with BSA PBS/0.3% Triton X-100/3% BSA for a minimum of 45 minutes. Incubation with TOMM20 (ABClonal, A6774) and IAV M1 (GeneTex, GTX40910) primary antibodies was performed in PBS overnight at 4°C. Cells were then washed three times in PBS and fluorescence staining was performed for one hour at room temperature using highly cross-adsorbed Alexa Fluor 488, 546 or 647 secondary antibodies at a concentration of 0.1mg/mL (Thermo Fisher Scientific). Following primary and secondary antibody incubations, cells were washed three times in PBS. To visualize nuclei, cells were stained with 1 μg/mL Hoechst 33342 in PBS for 10 minutes. Coverslips were mounted on microscope slides using ProLong Glass (ThermoFisher). Images were obtained using a DeltaVision Elite microscope system, Leica Thunder Imaging system, or a Zeiss LSM700 confocal microscope.

### Image Analysis, Quantification and Presentation

Image preparation and analysis was performed using Fiji (15). Deconvolved slices were collapsed into max intensity Z projections and brightness/contrast adjusted in Fiji before analysis.

Quantification of mitochondrial networks was performed using the ImageJ plugin, Mitochondria Analyzer (16). All experiments used the subtract background setting at 25 microns, enhance local contrast setting with max slope of 4, adjusted gamma setting at 1, and weighted mean method of thresholding. Block size and C-value was independently determined for each experiment using the 2D threshold optimize feature. All 37°C to 33°C image comparisons used the same brightness/contrast adjustment settings, threshold settings, and mitochondrial analyzer settings.

### Measuring Mitochondrial Membrane Potential

Cells were plated on µ-Slide 8 well imaging chambers for live cell imaging. The following day cells were treated with 25 µM Mdivi-1 or 500 µM Conezyme Q10 for 12 hours and then incubated in complete media containing MitoTracker Green (Thermo, M7514) and Tetramethylrhodamine, methyl ester (Thermo, I34361) for 30 minutes. Following incubation, cells were placed in complete media for imaging. Imaging was performed at 37°C in 5% CO2. Following acquisition, single cells were segmented from images and the MitoTracker Green channel was used to make a morphological mask of the mitochondria in Mitochondrial Analyzer. Next, Multiple Channel Analysis was used to obtain a ratio of the fluorescent signal from each channel for quantification.

### Tight Junction and Apoptosis Analysis and Apoptosis

MDCK-SIAT cells were plated on 24-well transwell inserts (Corning, 3470) at a density of 20,000 cells (37°C) or 60,000 cells (33°C) and incubated for 3-days prior to infection to reach confluency. Polarized cell cultures were then infected with IAV as described above. For tight junction analysis, cells were stained with antibodies to ZO-1 (Thermo, 40-2200) and IAV M1 (GeneTex, GTX40910) for 1 hour at room temperature to visualize tight junctions and viral antigen in mock infected and infected cells. Cells were then washed three times with PBS and stained for one hour at room temperature with previously mentioned secondary antibodies. ZO-1 signal was thresholded to create binary images from 500×500 pixel regions and quantified using Analyze > Measure > Area. For apoptosis analysis, Apopxin Green Indicator from the Apoptosis/Necrosis Assay Kit (ABCam, ab176749) was added to the apical portion of the transwell insert for 1 hour. The cells were then fixed and stained with IAV M1 (GeneTex, GTX40910). Apopxin Green signal was thresholded to create binary images and quantified using Analyze > Measure > Area.

### Oxygen Consumption Measurements

MDCK-SIAT cells were seeded onto an XF96e cell culture microplate at 30,000 cells/well and incubated overnight in complete media. On the day of measurement, cells were washed twice with Seahorse XF DMEM Basal Medium supplemented with 10 mM glucose, 2 mM l-glutamine, and 1 mM sodium pyruvate and preincubated for 1 h in a 37°C or 33°C humidified CO2-free incubator. OCR was measured by a Seahorse XF96e FluxAnalyzer with the Mito Stress Test kit (Agilent). The Mito Stress Test inhibitors were injected during the measurements as follows; oligomycin (2 μM), FCCP (1.0 μM), rotenone, and antimycin A (0.5 μM). The OCR values were normalized to cell density determined by the CyQUANT Cell Proliferation Assay Kit (Invitrogen, C7026). For OCR experiments with Mdivi-1, CoQ10, and vehicle treatments, the treatments were added 12 hours prior to analysis when analyzing treatment effect, or 12 prior to infection for infection experiments.

### Citrate Synthase Assay

The Citrate Synthase assay was performed using The Citrate Synthase Activity Assay kit (Sigma, MAK193). 1 x 10^6^ Cells were homogenized in 100 µL of ice-cold Citrate Synthase Assay Buffer Keep on ice for 10 minutes. Samples were then centrifuged at 10,000 x g for 5 minutes to remove insoluble material. 25 µL of sample was added to duplicate wells and brought to a volume of 50 µL using Citrate Synthase Assay Buffer. The plate was incubated at 25°C and A_412_ measurements were taken every 5 minutes for 35 minutes.

### Statistical analysis

All data were represented as mean ± SD. Data were analyzed in GraphPad Prism 10.0 software using Student’s t-test. Statistical significance was set at a threshold of P < 0.05. Data are representative of n = 3 biological replicates, unless stated otherwise in the figure legend.

## Results

### Human respiratory tract temperatures induce temperature-dependent alterations to the structure and function of mitochondrial networks

To evaluate temperature-dependent changes to mitochondrial networks, MDCK-SIAT cells were incubated at 33°C, a temperature consistent with the upper respiratory tract, or 37°C, a temperature consistent with the lower respiratory tract and evaluated using immunofluorescence microscopy. MDCK-SIAT cells are a commonly used cell line that is permissive to replication with human, clinical isolates of IAV such as the A/Baltimore/R0227/2017 (H3N2 clade 3c.2a2) used in this study. Mitochondrial networks in MDCK-SIAT cells incubated at 33°C contained a greater number of mitochondria and occupied a greater amount of area within the cell but displayed individual mitochondria that were less elongated when compared to mitochondrial networks in MDCK-SIAT cells incubated at 37°C (Fig 1A and 1C-E). Mitochondrial morphology is closely linked to functional properties of the organelle, with changes in shape and structure directly influencing cellular energy production and mitochondrial signaling networks (12, 17). To determine if respiratory tract temperatures influence functional properties of mitochondria in MDCK-SIAT cells, mitochondrial membrane potential was measured in cells at 33°C and 37°C using Tetramethylrhodamine, methyl ester (TMRM), a cationic cell-permeant dye that accumulates in a manner proportional to the degree of mitochondrial membrane potential.

**Figure 1:**
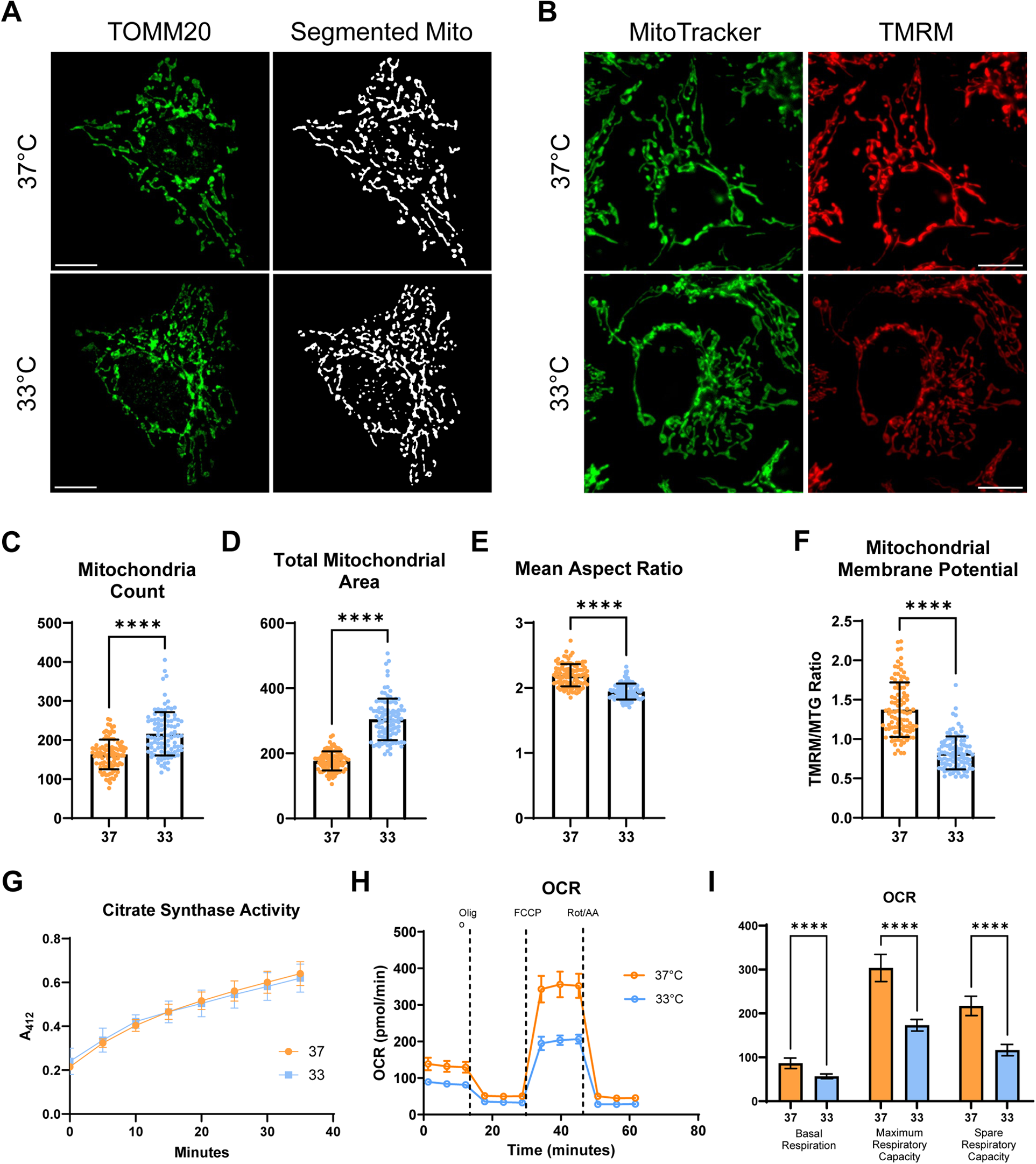
Incubating kidney epithelial cells at 33°C and 37°C induces temperature-specific structural and functional changes to mitochondrial networks. MDCK-SIAT cells incubated at 37°C or 33°C for 96 hours were **(A)** immunostained with anti-TOMM20 antibody or **(B)** co-stained with MitoTracker Green and TMRM. **(C-F)** Single cell images were used for thresholding and quantification of mitochondrial network properties using the imgaeJ plugin, Mitochondrial Analyzer (N = 100 cells at each temperature from three independent experiments). **(G)** Citrate synthase activity was measured from cell lysates using a spectrophotometric assay **(H-I)** Representative experiment of cellular oxygen consumption rate (OCR) measured in MDCK-SIAT cells incubated at 37°C or 33°C for 96 hours using a Seahorse XF96e FluxAnalyzer with the Mito Stress Test kit. Significant differences were determined by unpaired t test ****p < 0.0001.

TMRM was used in combination with MitoTracker Green (MTG), a thiol reactive live cell dye that labels mitochondria independent of mitochondrial function. The ratio of TMRM to MTG fluorescence was used as a measurement of membrane potential. Cells incubated at 33°C displayed lower mitochondrial membrane potential when compared to cells incubated at 37°C potentially indicating compromised mitochondrial functions like cellular respiration and ATP production in 33°C cells (Fig 1B and 1F). To further investigate the impact of temperature on metabolism a Seahorse XF96 Extracellular Flux Analyzer was used to measure the oxygen consumption rate (OCR) of cells at 33°C and 37°C. As expected, OCR measurements indicating basal respiration, maximum respiratory capacity and spare respiratory capacity were all significantly lower in cells incubated at 33°C compared to cells incubated at 37°C (Fig 1G-H). To better understand the source of the temperature-dependent differences in cellular metabolism citrate synthase activity was assessed, citrate synthase is the rate limiting enzyme of the citric acid cycle and its enzymatic activity can provide valuable information about mitochondrial content, overall mitochondrial function, metabolism, and mitochondrial biogenesis. No significant differences between 33°C and 37°C were found (Fig 1I) indicating that the increase in the overall number and area of mitochondria in cells incubated at 33°C may help compensate for the reduced respiratory capacity.

### IAV infection at 33 and 37°C produces different amounts of infectious virus results in temperature-specific changes to mitochondrial networks

IAV actively targets mitochondrial dynamics and mitochondrial signaling pathways during infection to mediate antiviral immune responses and aid virus replication (18–22). To investigate the impact of respiratory tract temperature on IAV replication and mitochondrial targeting, MDCK-SIAT cells incubated at 33°C and 37°C were infected with A/Baltimore/R0227/2017, an H3N2 (3c.2a2) clinical isolate from the 2017-18 influenza season. A high multiplicity of infection (MOI) and early timepoints post infection were chosen to study the temporal dynamics and viral host manipulation during a single cycle of replication. Cells incubated at 33°C and infected with IAV took longer to produce infectious virus (6 hours at 37°C vs 9 hours at 33°C) and produced significantly less infectious virus at 9 and 12 hours post infection than IAV infection of cells incubated at 37°C (Fig 2A). In addition to the differences in virus production, mitochondrial networks were evaluated at 6, 9, and 12 hours post infection at 33°C and 37°C to determine if there were unique, temperature-dependent changes to mitochondria during IAV infection. Notably, in accordance with previous studies performed at 37°C, elongation of mitochondria was observed at both temperatures during IAV and H1N1 infection (23). In addition to these observations, total mitochondrial area in infected cells at 37°C remained unchanged over the course of the 12-hour infection, while infected cells at 33°C showed a significant decrease in mitochondrial area per cell as early as 6 hours post infection (Fig 2B-C). The overall decrease in mitochondrial area was also observed in cells infected with an H1N1 clinical isolate from the 2019-20 influenza season (Fig S1). Interestingly, these results show that that underlying changes in host cell biology due to physiological range of temperature can directly influence infectious virus production and produce temperature-specific host cell outcomes during IAV infection.

**Figure 2:**
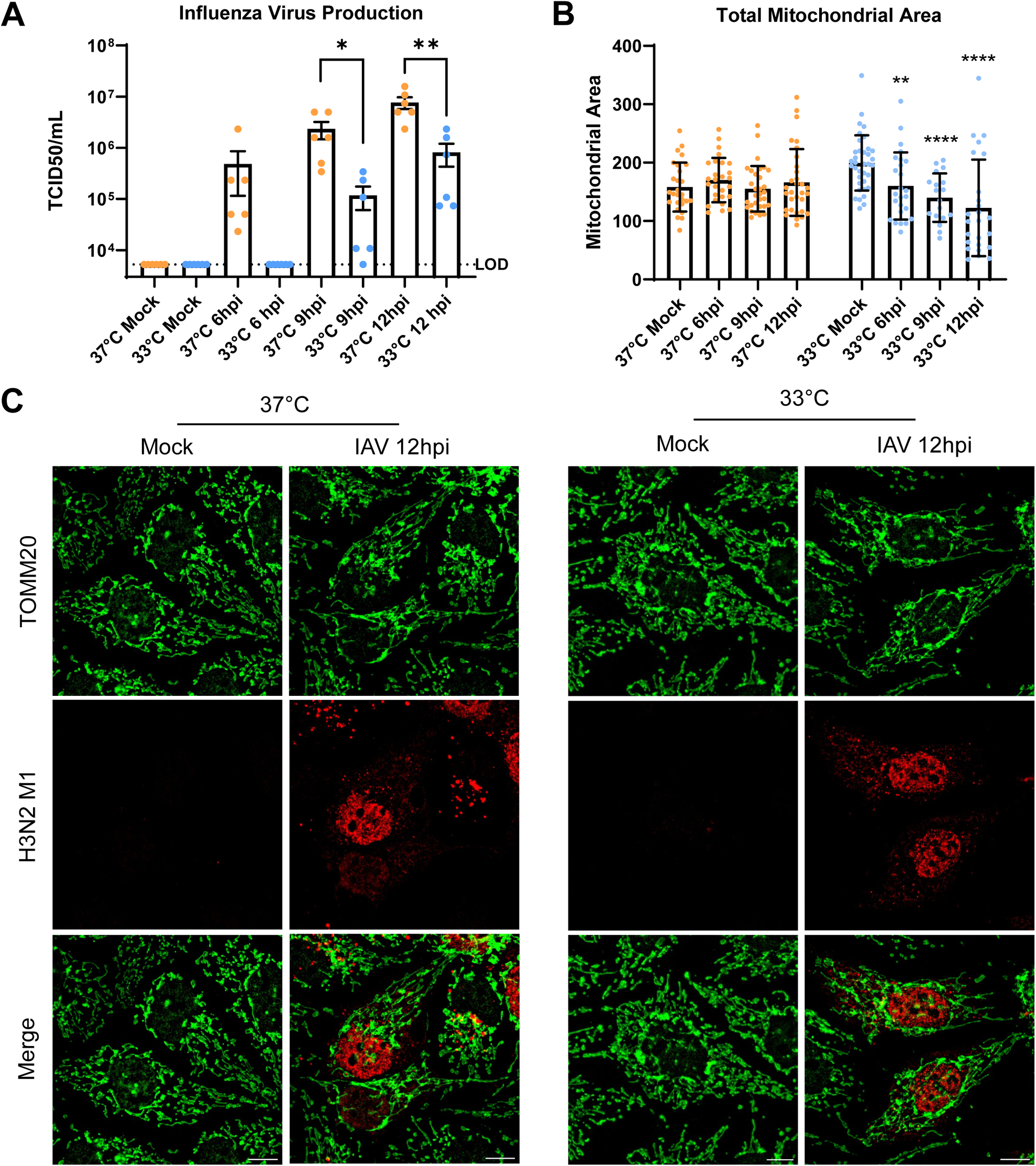
IAV infection at 33°C and 37°C produces different levels of infectious virus and temperature-specific changes to mitochondrial networks. **(A)** Infectious virus production at 37°C and 33°C from IAV infected MDCK-SIAT cells (MOI 5) quantified by TCID_50_ graph combines data from two independent experiments each with three technical replicates **(B)** Quantification of total area of mitochondrial networks in mock and IAV (MOI 5) infected cells at 6, 9, and 12 hours post infection. Each data point represents a single cell. Significance relative to mock infected cells at each temperature **(C)** Representative images of mock and IAV infected cells (MOI 5) at 12 hours post infection at 33°C and 37°C. Significant differences were determined by unpaired t test *p< 0.05 **p< 0.01 ****p < 0.0001.

### Respiratory tract temperature modulates the impact of IAV Infection on tight junctions, apoptosis induction, and host cell metabolism

Mitochondria play an important role in regulating effects of infection on host cells, impacting outcomes at both the cellular and tissue levels of organization. To gain a thorough understanding of host cell processes that could be impacted by changes in mitochondria due to respiratory tract temperature during infection measurements of tissue integrity, apoptosis, and cellular respiration during IAV infection of cells incubated at 33°C and 37°C were obtained. In the respiratory tract, tight junction integrity maintains barrier functions and helps to control inflammation and viral dissemination during virus infection (24, 25). Mitochondria help regulate tight junctions in MDCK cells by localizing to junctions and providing ATP needed for phosphorylation of several tight junction components (26, 27). Tight junction signal was measured over twenty-four hours of infection in polarized MDCK-SIAT cultures. Both 33°C and 37°C cell cultures displayed similar trends with a significant loss of tight junction signal at both 12 and 24 hours post infection (Fig 3D and S2) although the 33°C cultures, on average, lost a greater percentage of tight junction signal (40.91%) compared to 37°C cultures (30.17%) at 12 hours post infection.

**Figure 3:**
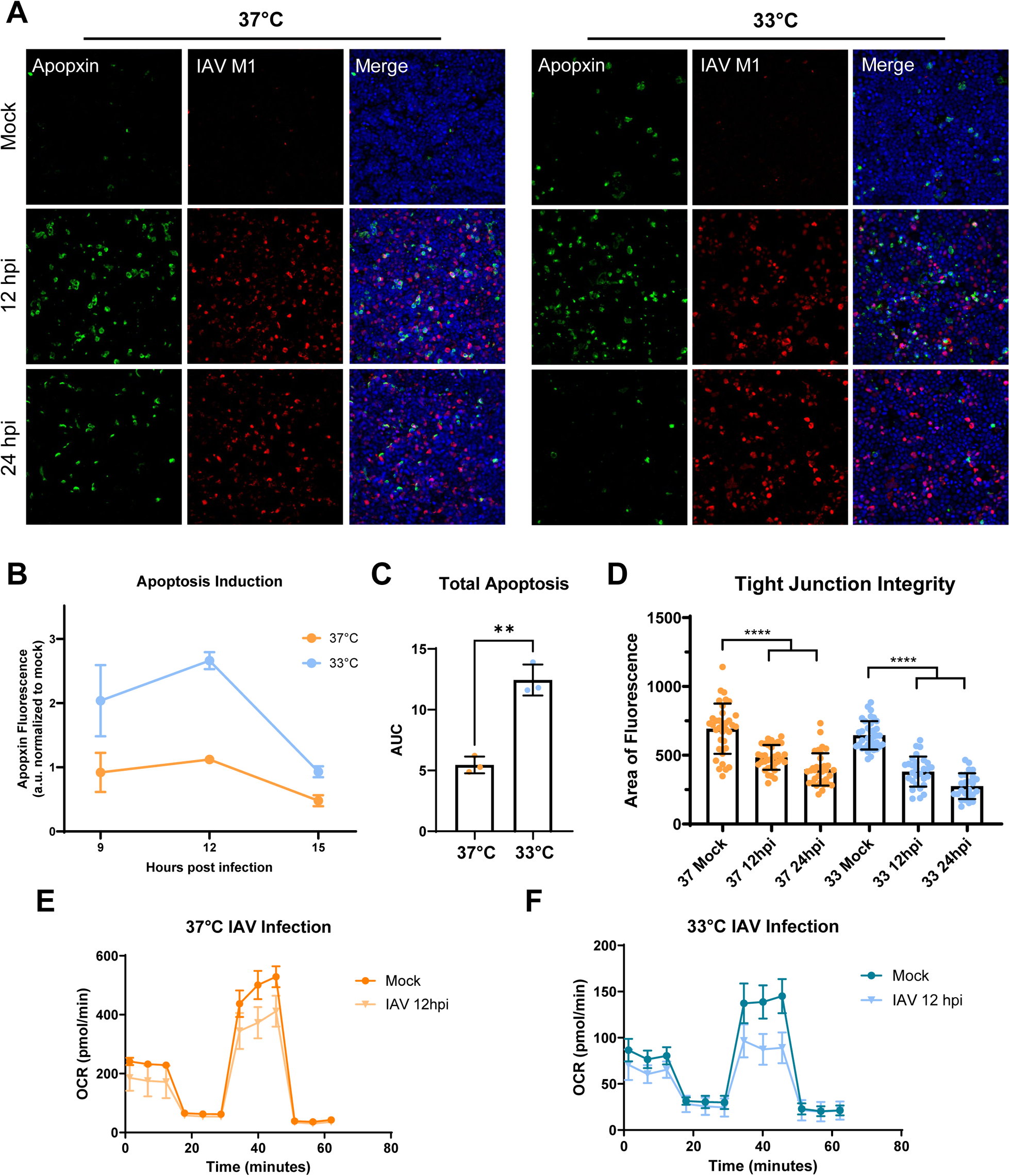
Physiological range of temperature modulates the impact of IAV Infection on tight junctions, apoptosis induction, and host cell metabolism. **(A)** 33°C and 37°C polarized MDCK-SIAT cells infected with IAV (MOI 5) and stained with Apopxin Green dye and anti-IAV M1 primary antibodies to detect apoptosis at 12 and 24 hours post infection. **(B)** Quantification of apoptosis at 9, 12, and 15 hours post infection from Apopxin Green fluorescence relative to mock infected transwells, graph represents combined replicates from 3 independent experiments. **(C)** Quantification of total apoptosis from 9-15 hours post infection using area under the curve **(D)** Quantification of tight junction signal from mock and IAV infected (MOI 5) polarized MDCK-SIAT cells using area of ZO-1 fluorescence graph represents data from two independent experiments with 25-35 infected regions analyzed per condition. **(E-F)** OCR measured in mock and IAV infected (MOI 5) MDCK-SIAT cells incubated at 37°C or 33°C for 12 hours post infection using a Seahorse XF96e FluxAnalyzer with the Mito Stress Test kit. Axis of graphs are adjusted to highlight the difference between mock and IAV infected conditions. Quantification of stress test curves present in Supplemental Figure 2, representative experiment shown from a total of two independent assays each with 33 technical replicates per condition. Significant differences were determined by unpaired t test ** P< 0.01 ****p < 0.0001.

Next, induction of apoptosis was analyzed in polarized MDCK-SIAT cultures, a process that has been implicated in the ability of IAV to regulate host immune responses with important implication in disease pathogenesis, and again found that the temporal trends between 33°C and 37°C cultures were similar. A significant increase in apoptosis was detected at 12 hours post infection followed by a decrease at 24 hours post infection at both temperatures (Fig 3A, S2D). However, cell cultures incubated at 33°C displayed a significantly higher amount of apoptosis at 12 hours post infection compared to the 37°C cultures (Fig S2D). To determine if this temperature specific difference was the result of altered apoptosis kinetics rather than a change in the overall magnitude of apoptosis, the kinetics of apoptosis induction were measured by obtaining measurements of cell death at 9, 12, and 15 hours post infection. IAV infected cells at 33°C exhibited greater levels of apoptosis than their counterparts at 37°C at all timepoints measured and displayed greater levels of apoptosis induction across the 9 - 15 hours post infection timespan (Fig 3C).

Lastly, the OCR of 33°C and 37°C cell cultures infected with IAV was measured at 12 hours post infection to determine the impact of infection on the metabolic state of the cell and to determine whether the baselines differences in respiratory capacity seen in Figure 1H-I would impact metabolic reprogramming during infection. Mitochondrial stress tests were performed on IAV infected cells 12 hours post infection and found significant decreases in the basal respiration, maximum respiratory capacity, and spare respiratory capacity at both 33°C and 37°C (Fig 3E-F, and S2B-C). Although both temperatures showed a significant reduction in respiration, the magnitude of reduction in maximal respiration was larger at 33°C (38.44%) than at 37°C (22.6%). Together, these results show that physiological range of temperatures can alter the magnitude and kinetics of a wide variety mitochondrial-related processes important for host cell outcomes during influenza infection.

### Mitochondrial modulating compounds can be used to recapitulate the morphological and functional differences observed in cells incubated at 33°C and 37°C

The temperature-dependent differences in cellular processes during IAV infection, particularly in processes tightly connected to mitochondrial dynamics, suggest that the temperature induced changes to mitochondrial networks play a direct role in the mediating these responses. To investigate this, the mitochondrial targeting small molecules, Mdivi-1 and Coenzyme Q10 were used to disentangle the specific contributions of mitochondria from other temperature-induced changes to cellular processes. Mdivi-1 inhibits the GTPase activity of the mitochondrial fission protein, Drp1, which results in the inhibition of mitochondrial fragmentation and the promotion of mitochondrial elongation. Coenzyme Q10 helps facilitate the transfer of electrons from complexes I and II to complex III in the electron transport chain, and exogenous supplementation has been previously shown to increase mitochondrial membrane potential and reduce reactive oxygen species. We hypothesized that treating cell cultures incubated at 33°C with these mitochondrial targeting small molecules would result in mitochondrial networks that mimic network properties seen in cells incubated at 37°C and would therefore result in viral replication kinetics and host cell outcomes that mimic those seen in cultures incubated at 37°C. Furthermore, given that these small molecules allow for the individual assessment of mitochondrial form and function each component could be evaluated to determine its contribution to the temperature-specific phenotypes observed in IAV infected cells. Mdvidi-1 treatment resulted in the elongation of mitochondria and increased mitochondrial area in cells cultured at both 33°C and 37°C (Fig 4A-B) without increasing mitochondrial membrane potential and Coenzyme Q10 (CoQ10) treatment increased mitochondrial membrane potential in MDCK-SIAT cells incubated at 33°C (Fig 4F) without altering the mitochondrial morphology.

**Figure 4:**
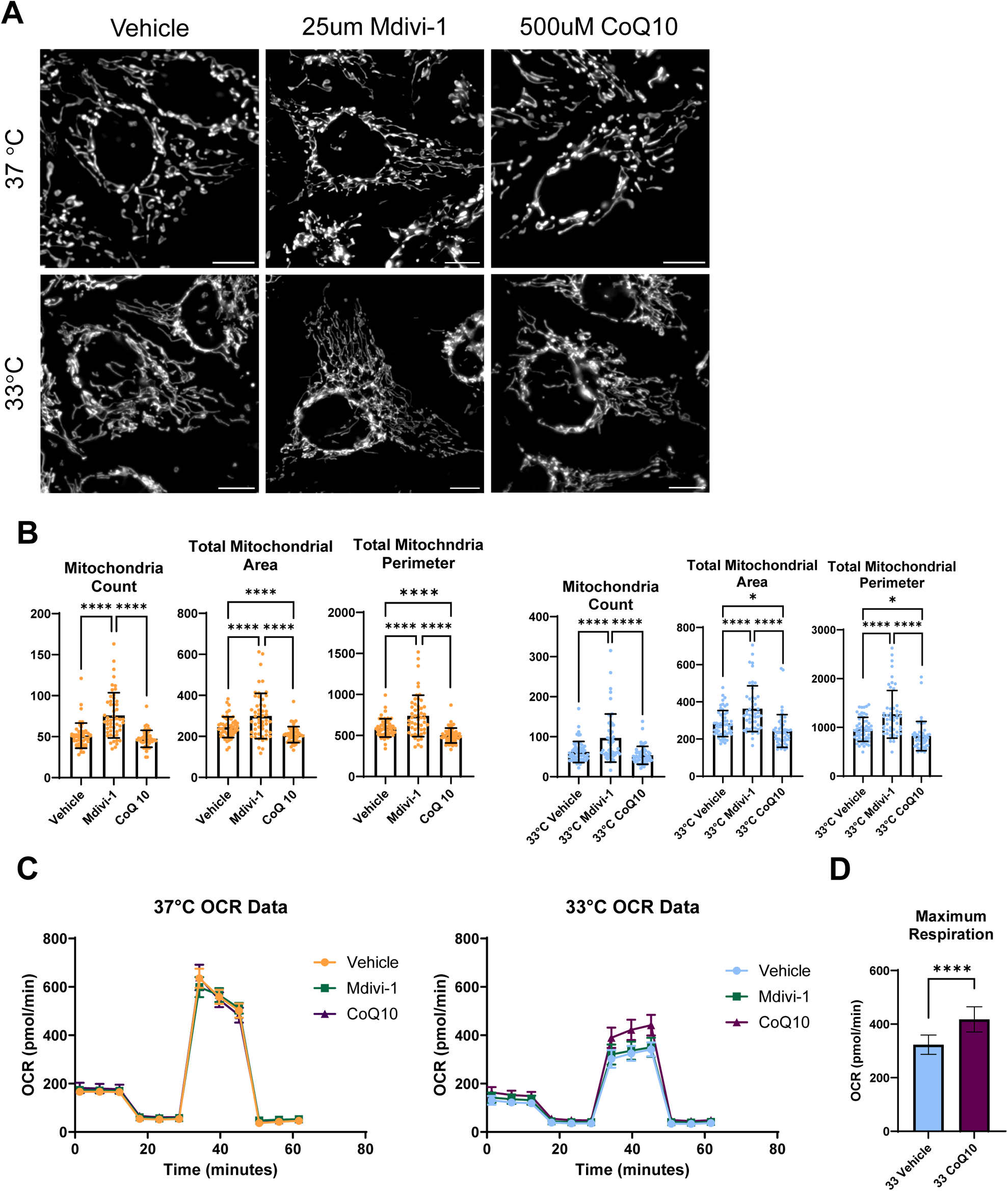
Mitochondrial modulating compounds Mdivi-1 and CoQ10 can be used to modulate morphological and functional properties of mitochondrial networks. **(A)** MDCK-SIAT cells incubated at 33°C and 37°C treated with Mdivi-1 and CoQ10 for 12 hours and stained with MitoTracker Green **(B)** Quantification of mitochondrial networks using Mitochondrial Analyzer graphs represent mitochondria analyzed from a minimum of 50 cells from two independent experiments. **(C)** OCR measured in MDCK-SIAT vehicle, Mdivi-1, and CoQ10 treated cells using a Seahorse XF96e FluxAnalyzer with the Mito Stress Test kit. A representative experiment from three independent replicates is shown. Significant differences were determined by unpaired t test * P< 0.05 ****p < 0.0001.

### Mitochondrial network modulators impact the dynamics of infectious IAV production and reverse temperature specific phenotypes during infection

The impact of Mdivi-1 and CoQ10 treatments on IAV infectious virus production was evaluated in MDCK-SIAT cells incubated at 33°C and 37°C and, as predicted, both Mdivi-1 and CoQ10 treatments resulted in the production of significantly greater amounts of virus over a 12 hour time period at both 33°C and 37°C indicating both mitochondrial elongation and increased mitochondrial membrane potential aid influenza virus production (Fig 5A). Next, the impact of IAV infection on host cell mitochondrial networks in Mdivi-1 and CoQ10 treatment cells was assessed using immunofluorescence microscopy to quantify mitochondrial area. We hypothesized that changes in mitochondrial networks from Mdivi-1 and CoQ10 treatments would cause mitochondria in cells incubated at 33°C to resemble mitochondrial networks in cells incubated at 37°C and therefore abrogate the temperature-dependent phenotypes observed during IAV infection. As anticipated, mitochondrial elongation via Mdivi-1 treatment was able to prevent the loss of mitochondrial mass at 12 hours post infection in IAV infected MDCK-SIAT cells incubated at 33°C, however, interestingly, CoQ10 treatment did not impact mitochondrial area during infection at 33°C or 37°C (Fig5B-F). Overall, these data indicate that the morphology of mitochondrial networks prior to infection, but not the functional capacity, can alter virus induced changes to mitochondrial networks during infection, and that changes in both the form and function of mitochondrial networks may play a direct role in mediating infectious virus production. Furthermore, these data support the conclusion that temperature-dependent differences in virus production may be directly mediated by temperature-dependent changes to mitochondrial networks.

**Figure 5:**
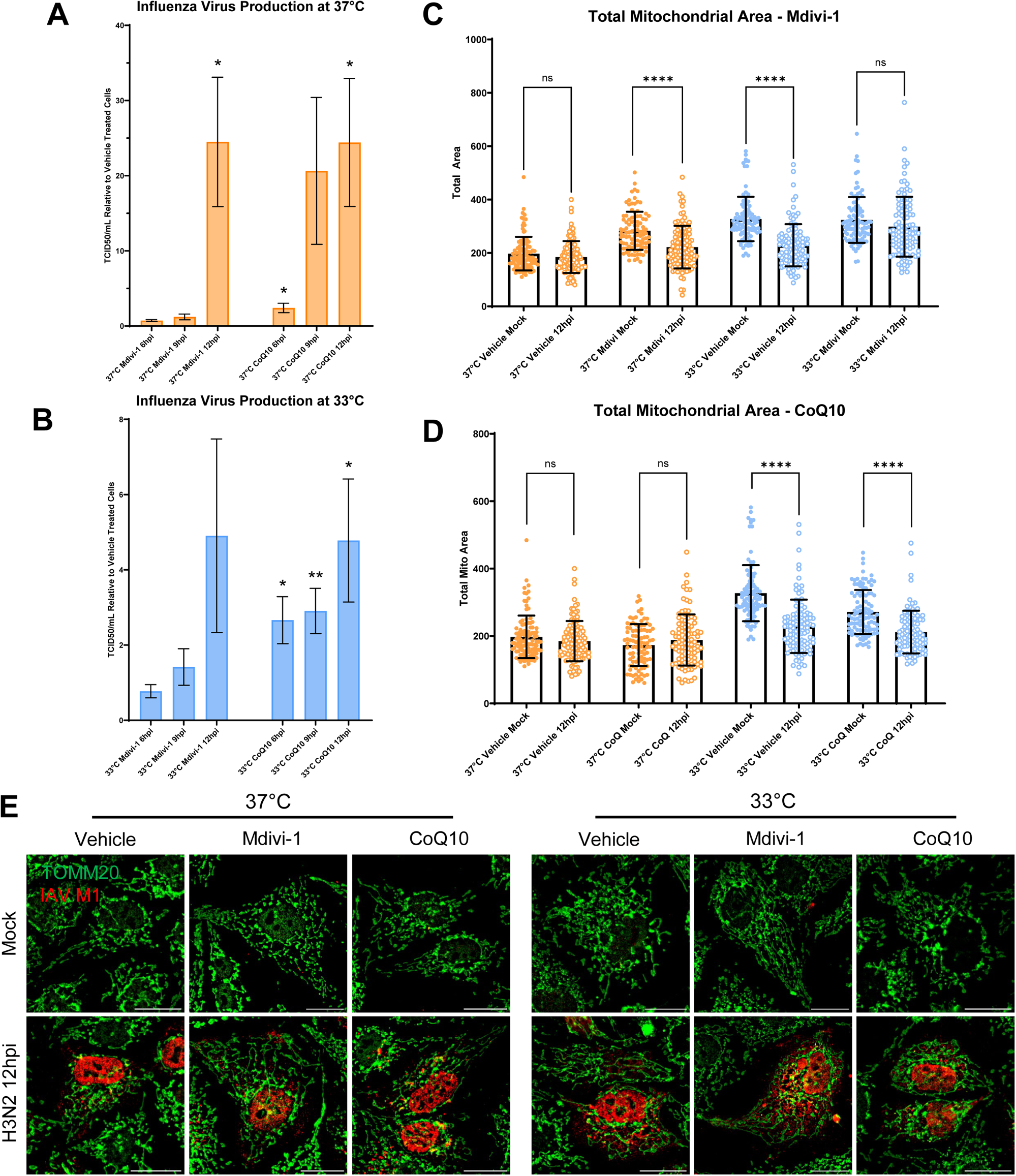
Mitochondrial network modulators impact the amount of infectious virus production and reverse temperature-specific phenotypes during infection. **(A-B)** Infectious IAV production at 37°C and 33°C in Mdivi-1 and CoQ10 treated MDCK-SIAT cells quantified by TCID_50_ graphed relative to vehicle treated cells at corresponding temperatures and timepoints. Graphs combine data from three independent experiments each with 4 technical replicates,significance is relative to vehicle treated cells at corresponding timepoints **(C-D)** Quantification of total area of mitochondrial networks in mock and IAV (MOI 5) vehicle, Mdivi-1, and CoQ10 treated cells at 6, 9, and 12 hours post infection. Each data point represents a single cell. Graphs represent data from a minimum of 100 cells collected from 3 independent experiments. **(E)** Representative images of mitochondrial networks from mock and IAV infected cells with mitochondrial treatments at 12 hours post infection at 33°C and 37°C. Significant differences were determined by unpaired t test * p< 0.05 ** p< 0.01 ****p < 0.0001.

### Upper respiratory tract temperature increases apoptosis induction during IAV infection independent of mitochondrial network modulators

To further investigate the extent to which specific, existing morphological and functional differences in mitochondrial networks impact host cell outcomes during IAV infection, changes to cellular respiration during IAV infection were evaluated in Mdivi-1 and CoQ10 treated cells. Across all treatment conditions and incubation temperatures IAV infection reduced the respiratory capacity of cells compared to mock infected cultures (Fig6A-D). Interestingly, despite increasing mitochondrial membrane potential in cells incubated at 33°C prior to infection, CoQ10 supplementation did not impact the loss of cellular respiration during infection or the magnitude of reduction in maximal respiration (35.8%) compared to vehicle treated 33°C cells (38.44%). This suggests that physiological temperature may be a more significant factor in determining the extent of cellular respiration loss during infection than the functional state of the mitochondria. Elongation of mitochondrial networks has previously been described to increase respiratory efficiency, however, Mdivi-1 treated cultures displayed lower basal respiration at baseline compared to vehicle treated cultures which may be due to off-target effects of Mdivi-1, which has been described to inhibit mitochondrial complex I in addition to the mitochondrial fission protein, Drp1. Regardless, mitochondrial elongation from Mdivi-1 treatment did not alter the decrease in cellular respiration due to IAV infection which provides further evidence that the loss of mitochondrial respiration during IAV infection occurs independent of mitochondrial functional capacity prior to infection. Altogether, these data indicate that shifts in cellular metabolism during IAV infection, characterized by a reduction in cellular respiration and an increase in glycolysis, occur independent of mitochondrial form or function and suggest that enhancing mitochondrial membrane potential alone is insufficient to counteract the impacts of IAV infection on cellular respiration.

**Figure 6:**
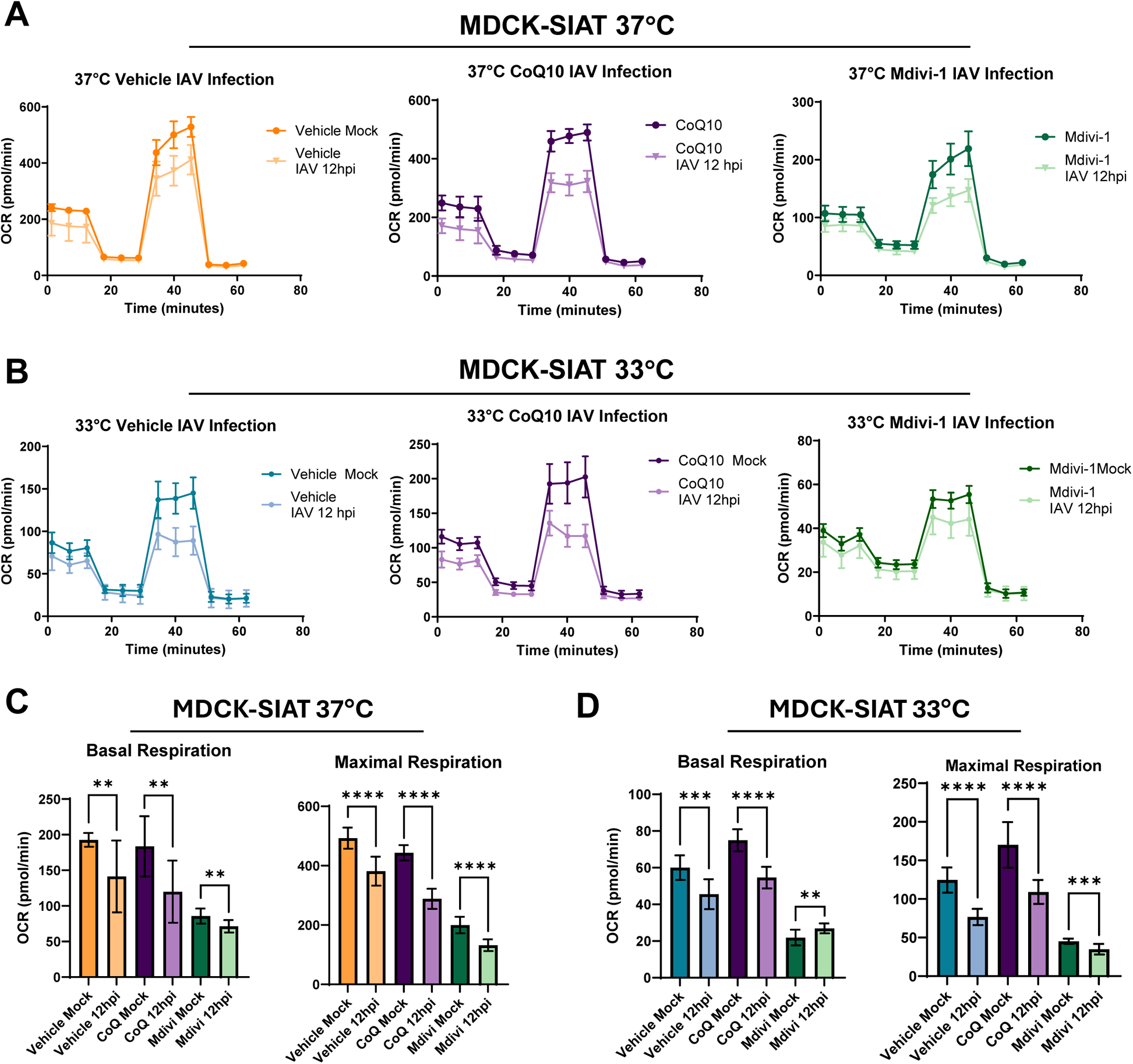
Modulating mitochondrial networks alone is insufficient to counteract the respiratory impairment caused by IAV infection. **(A-B)** Representative experiment of OCR measured 12 hours post infection in mock and IAV infected (MOI 5) MDCK-SIAT cells treated with vehicle, Mdivi-1 or CoQ10 at 37°C or 33°C. OCR measurements were collected using a Seahorse XF96e FluxAnalyzer with the Mito Stress Test kit. Axis of graphs are adjusted to highlight the difference between mock and IAV infected conditions. **(C-D)** Quantification of basal respiration and maximal respiration from panels A-B. Three independent experiments were conducted with 16 replicates per condition in each experiment. Significant differences were determined by unpaired t test ** P< 0.01 *** P< 0.001 ****p < 0.0001.

## Discussion

Physiological range of temperature has been shown to mediate host immune responses, virus production, and host cell outcomes, but the underlying changes to host cell biology at 33°C and 37°C mediating these differences are not understood. This study provides the first description of the physiological range of temperature modulating mitochondrial dynamics and explores how these changes impact host cell outcomes during IAV infection.

This study demonstrates that incubating MDCK cells at 33°C leads to an increase in the number of mitochondria and total mitochondrial area in the cell, but a decrease in the mitochondrial membrane potential and respiratory capacity when compared to cells at 37°C. In the context of IAV infection, we show that temperature influenced both the total amount of infectious virus produced during the first 12 hours of infection and the impact of viral infection on mitochondrial networks as measured by changes in total mitochondrial mass. Respiratory tract temperature has previously been shown to impact different aspects of IAV infection including replication kinetics and total virus production although the effects vary depending on the type of host cells and the genetic background of influenza virus (28–30). While we did not specifically investigate the mechanisms driving the loss of mitochondrial area during infection at 33°C, the bulk degradation of mitochondria via mitophagy is a likely pathway. The impacts of IAV infection on autophagy have been widely investigated and remain complicated as autophagy appears to be both an effective host defense for reducing viral antigen during infection while also aiding IAV replication and reducing host cell apoptosis (31). Interestingly, recent work has described an IAV nucleoprotein-mediated mechanism of mitophagy induction during infection which promotes IAV replication by inhibition of MAVS-mediated antiviral signaling (32, 33). One potential mechanism of mitochondrial loss in cells incubated at 33°C involves the degradation of the fragmented, low membrane potential mitochondria observed in 33°C cells via mitophagy to aid viral replication. This is supported by our findings that show only Mdivi-1-mediated elongation of mitochondria, which has been previously shown to protect against mitophagy (34), is able to rescue the loss of mitochondrial area during infection. Interestingly, the loss of mitochondrial area during infection does not appear to be coupled with infectious virus production giving support to previous hypotheses that the benefit of mitophagy induction to IAV replication centers around impaired innate immune signaling rather than nutrient recycling and shows that structural properties of mitochondrial networks play a role in modulating the ability of the virus to manipulate certain host cell dynamics.

Mitochondria are also known to play an important role in many processes relevant to host cell outcomes during IAV infection including metabolic programming, immune signaling, and apoptosis. In fact, previous studies have shown that the magnitude and temporal dynamics of mitochondrial dysfunction during viral infection can explain differences in disease severity between different types of respiratory virus infections, namely SARS-CoV-2 and IAV (35). It is also appreciated that mitochondrial morphodynamics greatly influence the cellular response after induction of these pathways. Therefore, given the underlying differences in mitochondrial networks observed in cells incubated at 33°C and 37°C a subset of these processes were evaluated and found to be impacted by respiratory tract temperature differences. One of the most notable results was the increased levels of apoptosis at 9 and 12 hours post infection in 33°C cell cultures compared to infected cultures at 37°C. Previous studies have shown that IAV infection induces intrinsic apoptosis via nucleoprotein expression and PB1-F2 interactions with mitochondrial proteins ANT3 and VDAC1 (10, 36–38). Although PB1-F2-induced apoptosis has not been shown to impact virus replication it has been implicated in the ability of the virus to successfully evade host immune responses and contribute to pathogenicity of infection (39). Few studies have evaluated the importance of apoptosis kinetics during IAV infection, and further research is needed to determine which aspects of the viral life cycle benefit from the prevention or induction of host cell apoptosis and whether IAV actively regulates the balance of apoptosis induction during infection. This research supports the idea that apoptosis dynamics may differ depending on the physiological ranges of temperature. Further exploring the role of apoptosis induction and its contribution to efficient evasion of immune cells in the context of disease pathogenesis will help provide context to the findings described here.

Despite changes in mitochondrial networks at 33°C and 37°C, not all mitochondrial related functions that were examined exhibited drastic temperature dependent phenotypes, most notably, IAV infected cells exhibited a reduction in cellular respiration regardless of incubation temperature or treatment conditions indicating that some mitochondrial-related processes may be impacted during infection independent of mitochondrial form and function. These changes may be so integral to IAV replication that few alterations in host cell biology can affect the virus’s ability to manipulate them. Of course, cellular respiration is only one component of metabolism and further research is needed to fully explore the impact of respiratory tract temperature on host cell metabolism. Previous work has shown increases in c-Myc, glycolysis, and glutaminolysis in IAV infected primary human respiratory cells which highlights additional areas to assess regarding the effects of temperature on metabolism (40). Additionally, the metabolic demands of activated immune cells that are recruited to sites of infection result in a competition for metabolic pathway intermediates which further highlights the importance of cell death and the transfer of nutrients as processes that have the potential to impact immune responses during IAV infection (9). The data presented here show the consistent loss of oxidative capacity in cells infected with IAV regardless of respiratory tract temperature, starting respiratory capacity, or mitochondrial manipulation with Mdivi-1 or CoQ10. These findings hint towards a central role for IAV-mediated manipulation of host cell metabolism during infection and provide a basis to expand metabolic studies in the context of respiratory tract temperature and viral infection.

Together, the data show that respiratory tract temperatures can lead to definitive changes in host cell mitochondrial biology, influence IAV replication, and impact host cell outcomes. These findings provide the first descriptions of how respiratory tract temperature can influence host cell dynamics and how these underlying changes can impact outcomes of IAV infection. Lastly, further research is needed to characterize the effect of respiratory tract temperature on primary, differentiated respiratory epithelial cells, focusing both on baseline biological differences at 33°C and 37°C across various epithelial cell types and differences during respiratory virus infection. Such cell culture models add physiological relevance to temperature studies and offer a promising path to uncover key cellular processes and functional aspects of the respiratory epithelium that are affected by temperature.

## Acknowledgements

This research reported in this publication was supported by the National Institute of Allergy and Infectious Diseases (NIAID), National Institutes of Health (NIH), Centers of Excellence for Influenza Research and Response (CEIRR) contract 75N93021C00045, the Richard Eliasberg Family Foundation, the Office of the Director of the National Institutes of Health under award number S10OD016374. We thank the laboratories of Nicole Baumgarth, Kimberly Davis, Sabra Klein and Andrew Pekosz for comments on and discussions about the data presented.

---

**Supplemental Figure 1:**
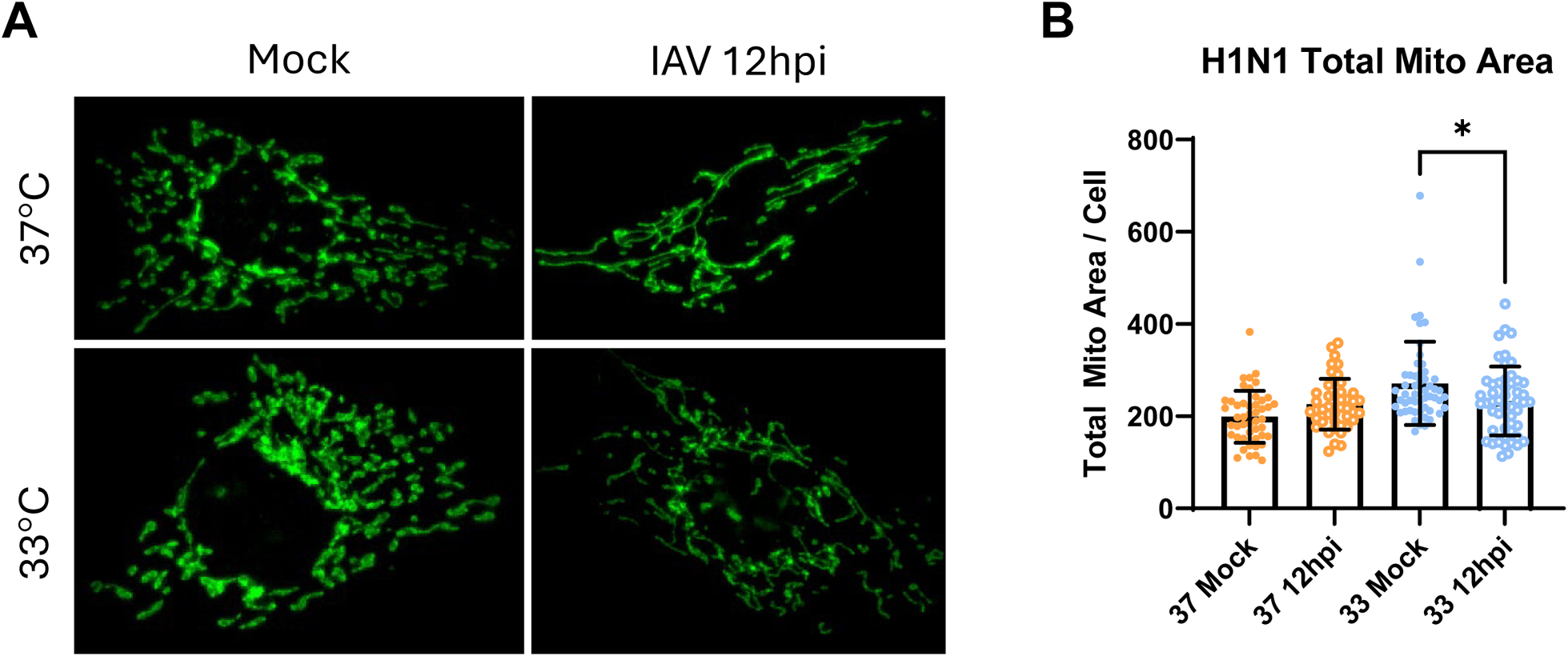
H1N1 IAV virus infection at 33°C and 37°C induces temperature-specific changes to mitochondrial networks. **(A)** Representative images of mock and H1N1 infected cells (MOI 5) at 12 hours post infection at 33°C and 37°C. **(B)** Quantification of total area of mitochondrial networks in mock and H1N1 (MOI 5) infected cells at 12 hours post infection. Each data point represents a single cell. Significant differences were determined by unpaired t test *p< 0.05

**Supplemental Figure 2:**
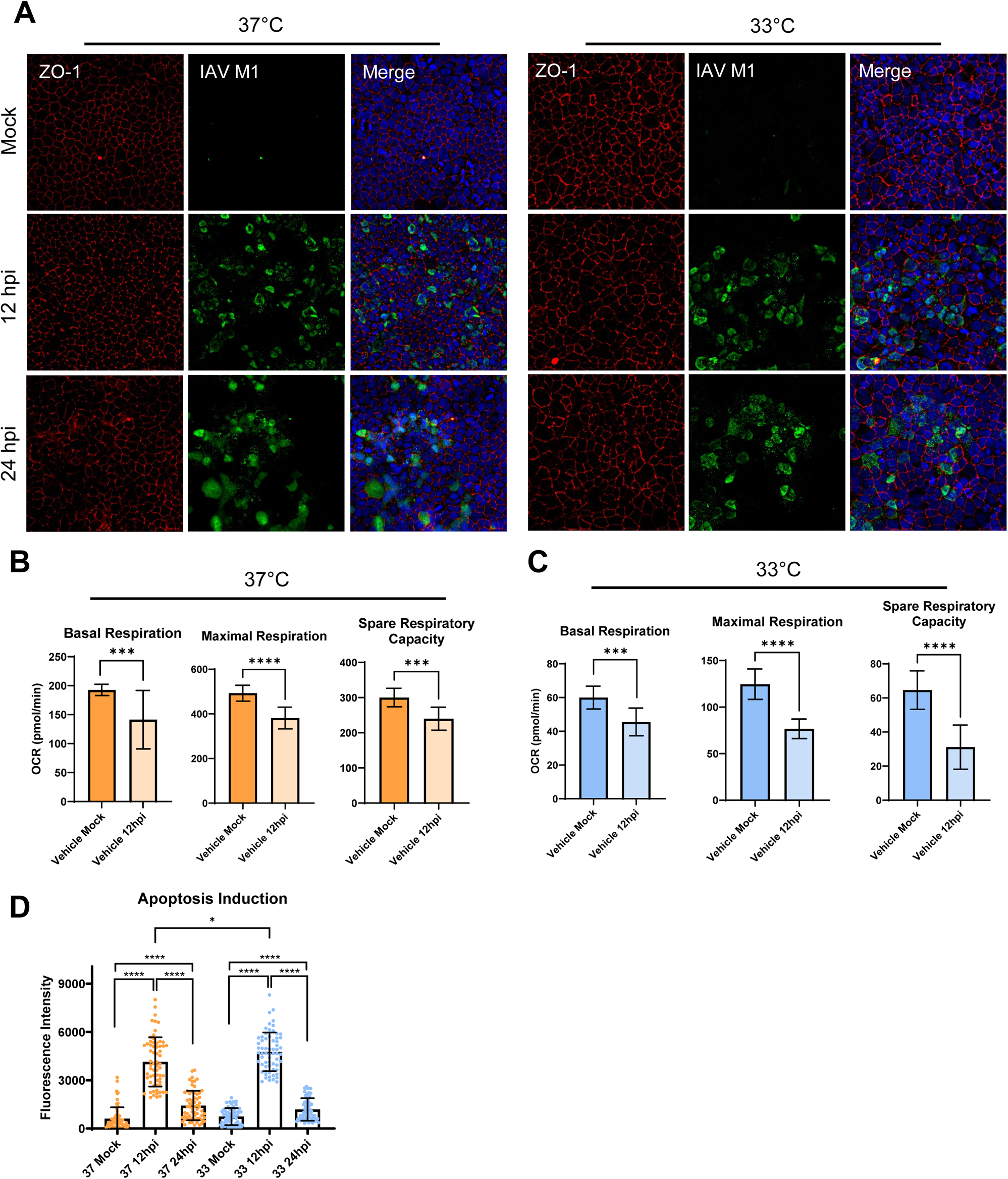
Physiological range of temperature modulates the impact of IAV Infection on tight junctions, apoptosis induction, and host cell metabolism. **(A)** 33°C and 37°C polarized MDCK-SIAT cells infected with IAV (MOI 5) and stained with anti-ZO-1 and anti-IAV M1 primary antibodies to detect tight junction degradation at 12 and 24 hours post infection. **(B)** Quantification of Mito Stress Tests performed on IAV infected (MOI 5) MDCK-SIAT cells incubated at 37°C or 33°C for 12 hours post infection **(C)** Quantification of Apopxin Green fluorescenece from IAV infected (MOI 5) polarized MDCK-SIAT cells at 12 and 24 hours post infection. Graphs combine data from two independent experiments. Significant differences were determined by unpaired t test ** P< 0.01 ****p < 0.0001.

**Supplemental Figure 3:**
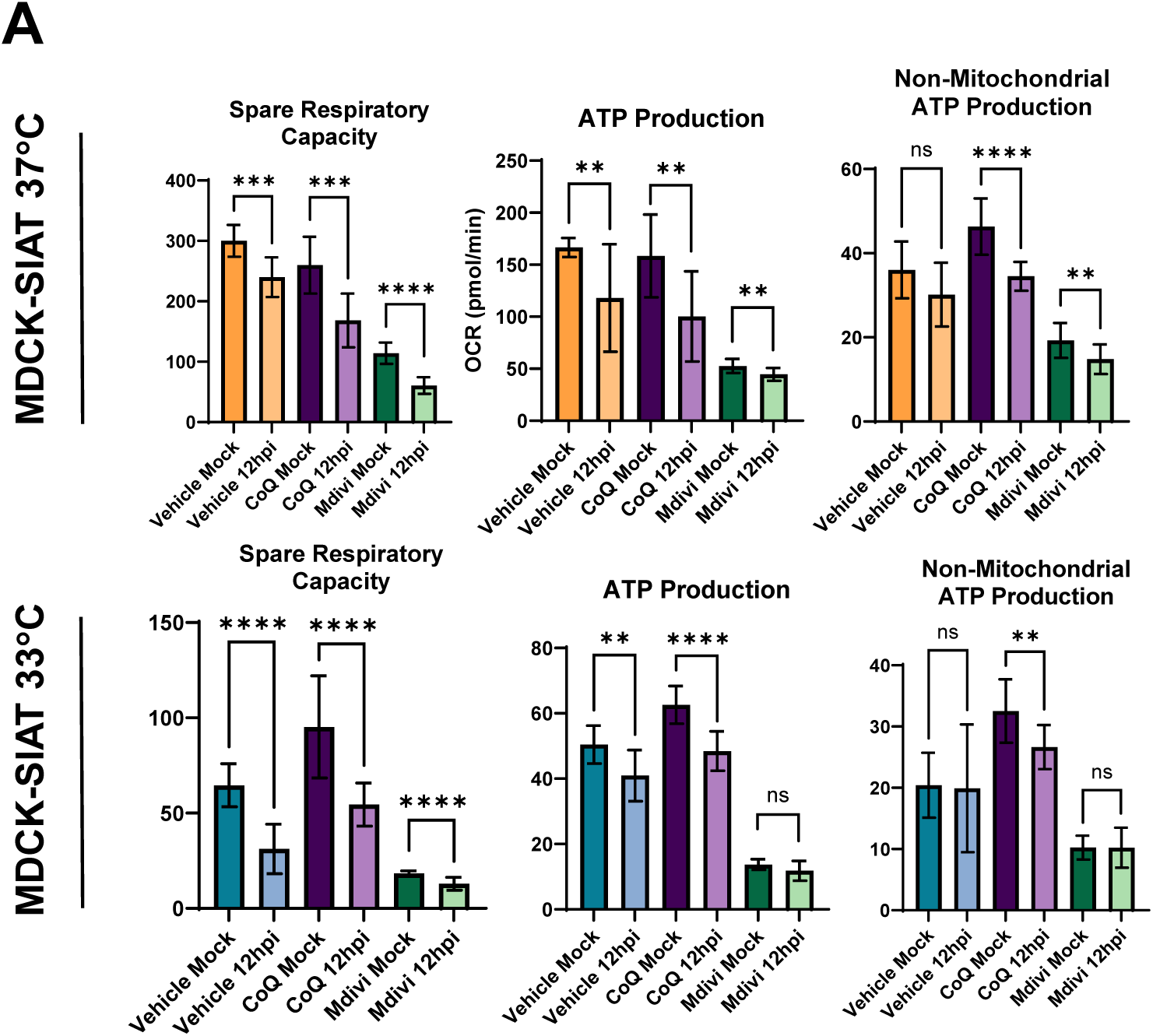
Modulating mitochondrial networks alone is insufficient to counteract the respiratory impairment caused by IAV infection. **(A)** Quantification from a representative Mito Stress Test performed 12 hours post infection on mock and IAV infected (MOI 5) MDCK-SIAT cells treated with vehicle, Mdivi-1 or CoQ10 at 37°C or 33°C. Three independent experiments were conducted with 16 replicates per condition in each experiment. Significant differences were determined by unpaired t test ** P< 0.01 *** P< 0.001 ****p < 0.0001.

## References

1. McFadden ER, Jr. Heat and water exchange in human airways. Am Rev Respir Dis. 1992;146(5 Pt 2):S8–10.

2. Keck T, Leiacker R, Riechelmann H, Rettinger G. Temperature profile in the nasal cavity. Laryngoscope. 2000;110(4):651–4.

3. Keck T, Leiacker R, Heinrich A, Kühnemann S, Rettinger G. Humidity and temperature profile in the nasal cavity. Rhinology. 2000;38(4):167–71.

4. Peteranderl C, Herold S, Schmoldt C. Human Influenza Virus Infections. Semin Respir Crit Care Med. 2016;37(4):487–500.

5. Foxman EF, Storer JA, Fitzgerald ME, Wasik BR, Hou L, Zhao H, et al. Temperature-dependent innate defense against the common cold virus limits viral replication at warm temperature in mouse airway cells. Proc Natl Acad Sci U S A. 2015;112(3):827–32.

6. Resnick JD, Beer MA, Pekosz A. Early Transcriptional Responses of Human Nasal Epithelial Cells to Infection with Influenza A and SARS-CoV-2 Virus Differ and Are Influenced by Physiological Temperature. Pathogens. 2023;12(3).

7. V’Kovski P, Gultom M, Kelly JN, Steiner S, Russeil J, Mangeat B, et al. Disparate temperature-dependent virus-host dynamics for SARS-CoV-2 and SARS-CoV in the human respiratory epithelium. PLoS Biol. 2021;19(3):e3001158.

8. Iwasaki A, Pillai PS. Innate immunity to influenza virus infection. Nat Rev Immunol. 2014;14(5):315–28.

9. Bahadoran A, Bezavada L, Smallwood HS. Fueling influenza and the immune response: Implications for metabolic reprogramming during influenza infection and immunometabolism. Immunol Rev. 2020;295(1):140–66.

10. Gui R, Chen Q. Molecular Events Involved in Influenza A Virus-Induced Cell Death. Front Microbiol. 2021;12:797789.

11. Tait SW, Green DR. Mitochondria and cell signalling. J Cell Sci. 2012;125(Pt 4):807–15.

12. Picard M, Shirihai OS, Gentil BJ, Burelle Y. Mitochondrial morphology transitions and functions: implications for retrograde signaling? Am J Physiol Regul Integr Comp Physiol. 2013;304(6):R393–406.

13. Reed LJ, Muench H. A SIMPLE METHOD OF ESTIMATING FIFTY PER CENT ENDPOINTS12. American Journal of Epidemiology. 1938;27(3):493–7.

14. McCown MF, Pekosz A. The influenza A virus M2 cytoplasmic tail is required for infectious virus production and efficient genome packaging. J Virol. 2005;79(6):3595–605.

15. Schindelin J, Arganda-Carreras I, Frise E, Kaynig V, Longair M, Pietzsch T, et al. Fiji: an open-source platform for biological-image analysis. Nat Methods. 2012;9(7):676–82.

16. Chaudhry A, Shi R, Luciani DS. A pipeline for multidimensional confocal analysis of mitochondrial morphology, function, and dynamics in pancreatic β-cells. Am J Physiol Endocrinol Metab. 2020;318(2):E87–e101.

17. Glancy B, Kim Y, Katti P, Willingham TB. The Functional Impact of Mitochondrial Structure Across Subcellular Scales. Front Physiol. 2020;11:541040.

18. Sorouri M, Chang T, Hancks DC. Mitochondria and Viral Infection: Advances and Emerging Battlefronts. mBio. 2022;13(1):e0209621.

19. Elesela S, Lukacs NW. Role of Mitochondria in Viral Infections. Life (Basel). 2021;11(3).

20. Foo J, Bellot G, Pervaiz S, Alonso S. Mitochondria-mediated oxidative stress during viral infection. Trends Microbiol. 2022;30(7):679–92.

21. Purandare N, Ghosalkar E, Grossman LI, Aras S. Mitochondrial Oxidative Phosphorylation in Viral Infections. Viruses. 2023;15(12).

22. Saxena R, Sharma P, Kumar S, Agrawal N, Sharma SK, Awasthi A. Modulation of mitochondria by viral proteins. Life Sci. 2023;313:121271.

23. Pila-Castellanos I, Molino D, McKellar J, Lines L, Da Graca J, Tauziet M, et al. Mitochondrial morphodynamics alteration induced by influenza virus infection as a new antiviral strategy. PLoS Pathog. 2021;17(2):e1009340.

24. Guttman JA, Finlay BB. Tight junctions as targets of infectious agents. Biochim Biophys Acta. 2009;1788(4):832–41.

25. Linfield DT, Raduka A, Aghapour M, Rezaee F. Airway tight junctions as targets of viral infections. Tissue Barriers. 2021;9(2):1883965.

26. Madan S, Uttekar B, Chowdhary S, Rikhy R. Mitochondria Lead the Way: Mitochondrial Dynamics and Function in Cellular Movements in Development and Disease. Front Cell Dev Biol. 2021;9:781933.

27. Tsukamoto T, Nigam SK. Role of tyrosine phosphorylation in the reassembly of occludin and other tight junction proteins. Am J Physiol. 1999;276(5):F737–50.

28. Fischer WA, 2nd, King LS, Lane AP, Pekosz A. Restricted replication of the live attenuated influenza A virus vaccine during infection of primary differentiated human nasal epithelial cells. Vaccine. 2015;33(36):4495–504.

29. Massin P, Kuntz-Simon G, Barbezange C, Deblanc C, Oger A, Marquet-Blouin E, et al. Temperature sensitivity on growth and/or replication of H1N1, H1N2 and H3N2 influenza A viruses isolated from pigs and birds in mammalian cells. Vet Microbiol. 2010;142(3-4):232–41.

30. Yamaya M, Nishimura H, Lusamba Kalonji N, Deng X, Momma H, Shimotai Y, et al. Effects of high temperature on pandemic and seasonal human influenza viral replication and infection-induced damage in primary human tracheal epithelial cell cultures. Heliyon. 2019;5(2):e01149.

31. Zhang R, Chi X, Wang S, Qi B, Yu X, Chen JL. The regulation of autophagy by influenza A virus. Biomed Res Int. 2014;2014:498083.

32. Zhang B, Xu S, Liu M, Wei Y, Wang Q, Shen W, et al. The nucleoprotein of influenza A virus inhibits the innate immune response by inducing mitophagy. Autophagy. 2023;19(7):1916–33.

33. Wang R, Zhu Y, Ren C, Yang S, Tian S, Chen H, et al. Influenza A virus protein PB1-F2 impairs innate immunity by inducing mitophagy. Autophagy. 2021;17(2):496–511.

34. Gomes LC, Di Benedetto G, Scorrano L. During autophagy mitochondria elongate, are spared from degradation and sustain cell viability. Nat Cell Biol. 2011;13(5):589–98.

35. Yeung-Luk BH, Narayanan GA, Ghosh B, Wally A, Lee E, Mokaya M, et al. SARS-CoV-2 infection alters mitochondrial and cytoskeletal function in human respiratory epithelial cells mediated by expression of spike protein. mBio. 2023;14(4):e0082023.

36. Chen W, Calvo PA, Malide D, Gibbs J, Schubert U, Bacik I, et al. A novel influenza A virus mitochondrial protein that induces cell death. Nat Med. 2001;7(12):1306–12.

37. Zamarin D, García-Sastre A, Xiao X, Wang R, Palese P. Influenza virus PB1-F2 protein induces cell death through mitochondrial ANT3 and VDAC1. PLoS Pathog. 2005;1(1):e4.

38. Tripathi S, Batra J, Cao W, Sharma K, Patel JR, Ranjan P, et al. Influenza A virus nucleoprotein induces apoptosis in human airway epithelial cells: implications of a novel interaction between nucleoprotein and host protein Clusterin. Cell Death Dis. 2013;4(3):e562.

39. Zamarin D, Ortigoza MB, Palese P. Influenza A virus PB1-F2 protein contributes to viral pathogenesis in mice. J Virol. 2006;80(16):7976–83.

40. Smallwood HS, Duan S, Morfouace M, Rezinciuc S, Shulkin BL, Shelat A, et al. Targeting Metabolic Reprogramming by Influenza Infection for Therapeutic Intervention. Cell Rep. 2017;19(8):1640–53.

